# Options for calibrating CERES-maize genotype specific parameters under data-scarce environments

**DOI:** 10.1101/353045

**Authors:** A.A Adnan, J. Diels, J.M. Jibrin, A.Y. Kamara, P. Craufurd, I. B. Mohammed, Z.E.H. Tonnang

**Author notes:** Current address: Department of Earth and Environmental Sciences, Division of Soil and Water Management, KU Leuven, Celestijnenlaan 200E, 3001 Leuven, Belgium.

## Abstract

Most crop simulation models require the use of Genotype Specific Parameters (GSPs) which provide the Genotype component of G×E×M interactions. Estimation of GSPs is the most difficult aspect of most modelling exercises because it requires expensive and time-consuming field experiments. GSPs could also be estimated using multi-year and multi locational data from breeder evaluation experiments. This research was set up with the following objectives: i) to determine GSPs of 10 newly released maize varieties for the Nigerian Savannas using data from both calibration experiments and by using existing data from breeder varietal evaluation trials; ii) to compare the accuracy of the GSPs generated using experimental and breeder data; and iii) to evaluate CERES-Maize model to simulate grain and tissue nitrogen contents. For experimental evaluation, 8 different experiments were conducted during the rainy and dry seasons of 2016 across the Nigerian Savanna. Breeder evaluation data was also collected for 2 years and 7 locations. The calibrated GSPs were evaluated using data from a 4 year experiment conducted under varying nitrogen rates (0, 60 and 120kg N ha^−1^). For the model calibration using experimental data, calculated model efficiency (EF) values ranged between 0.86-0.92 and coefficient of determination (d-index) between 0.92-0.98. Calibration of time-series data produced nRMSE below 7% while all prediction deviations were below 10% of the mean. For breeder experiments, EF (0.52-0.81) and d-index (0.46-0.83) ranges were lower. Prediction deviations were below 17% of the means for all measured variables. Model evaluation using both experimental and breeder trials resulted in good agreement (low RMSE, high EF and d-index values) between observed and simulated grain yields, and tissue and grain nitrogen contents. We conclude that higher calibration accuracy of CERES-Maize model is achieved from detailed experiments. If unavailable, data from breeder experimental trials collected from many locations and planting dates can be used with lower but acceptable accuracy.

## 1.0 Introduction

Maize has become an important crop in Nigeria in the past decades due to its importance as food for human consumption; feed for animals and also as a source of industrial raw material [1]. Despite its importance, yield of maize has remained quite low in the Savannas mostly due to biotic and abiotic constraints [2]. In recent years, new early and extra early maturing maize varieties that are tolerant to most of the biotic and abiotic constraints have been developed for the Nigerian Savannas by the International Institute for Tropical Agriculture (IITA) and its partners. Several agronomic technologies have also been developed to increase the productivity of these varieties with a view to enhancing maize productivity. Before the varieties are released, they are usually grown under multi-locational yield and crop management evaluation trials over several years. Dissemination of such varieties and technologies will require setting up of costly and time-consuming experiments across wide areas. This is needed in order to adequately evaluate the Genotype x Environment interaction which demonstrates the performance of each variety across diverse environments. Unless this is done, breeders cannot conclusively recommend genotypes for particular environments.

Crop simulation modeling offers an opportunity to explore the potential of new varieties and crop management practices in different environments (soil, climate, management) prior to their release [3]. Recently, use of crop simulation models, particularly DSSAT, is on the increase in Africa through initiatives such as the Agricultural Models Inter-Comparison Project (AgMIP) [4]. In West Africa, the CERES-Maize model has been recently used by McCarthy et al. [3] to evaluate climate-sensitive farm management practices in the Northern Regions of Ghana. Adnan et al. [5 and 6] used the same model to determine the nitrogen fertilization requirements of early maturing maize in the Sudan Savanna of Nigeria and the optimum planting dates of maize in Northern Nigeria. One of the major requirements for the use of crop simulations is calibration of Genotype Specific Parameters (GSPs). GSPs are sets of parameters that enable crop models to simulate the performance of diverse genotypes under varying soil, weather and management conditions [7]. Like all other parameters in crop simulation models, the GSPs must have a physical or biological meaning [8]. Measuring GSPs directly from real systems (farm and field level) is very complex and impractical, and results in highly inaccurate and uncertain values of estimated variables [9 and 10]. Direct measurement requires setting up of field or growth chamber studies, collection of many samples, and exposure to different photoperiods where necessary. The most common method for deriving GSPs is from field experiments designed specifically for their estimation [11 and 12]. This process is quite expensive, time consuming and requires regular sampling of growth, phenology and yield data for each variety following a set of minimum dataset rules [7].

Since most models have been developed elsewhere in Europe and USA, their use outside their domain of development requires a great deal of data for their calibration and validation. Several approaches for estimating GSPs have been documented. The genetic coefficient calculator (GENCALC) was used by Anothai et al. [13] to determine variety coefficients for new peanut lines in Thailand from standard varietal trials. From their experiments, they were able to successfully calibrate groundnut GSPs using a set of field experiments and yield evaluation experiments using the GENCALC software. Mavromatis et al. [14] successfully generated GSPs of soybean from crop performance trials in Georgia, USA. Bannayan et al. [15] employed a pattern recognition technique, which is based on similarity measures, to estimate GSPs for maize. In their experiments, pattern recognition was used as an alternative to GENCALC and GLUE in estimation of maize GSPs. The generalized likelihood uncertainty estimation (GLUE) method was used by He et al. [16] to successfully estimate maize GSPs in North Carolina.

With a growing number of researchers using the DSSAT model in the Savannas of Africa, there is need to evaluate the GSP calibration step as it is the aspect that requires the greatest amount of data and expertise. Calibration of GSPs can also be done using secondary data from breeders who routinely conduct multi-location trials. Such datasets are available in Africa where strong breeding programs are present. Because the conventional method of calibrating GSPs is quite expensive and laborious, there is need to utilize secondary breeder trial data for calibrating maize GSPs and to evaluate the accuracy of this approach by comparing it with calibrations done using detailed calibration experiments.

The objectives of this research were: i) to determine GSPs of 10 newly released open pollinated (OPV) and hybrid maize varieties for the Nigerian Savannas using data from both field experiments specifically designed for this purpose (herein called calibration experiments) and by using data from breeder varietal evaluation trials (herein called breeder evaluation experiments); ii) to compare the accuracy of the GSPs generated using calibration and breeder data; and iii) to evaluate the ability of the GSPs calibrated using the 2 methods to simulate grain yield and tissue/grain nitrogen contents of maize.

## 2.0 Materials and methods

### 2.1 Model description

The maize model used in this study is the CSM CERES-Maize model of DSSAT version 4.6. Detailed description of the CERES-maize model of DSSAT can be found in Jones et al [17]. CERES-maize is variety and site specific, and operates on a daily time step. It dynamically simulates the development of roots and shoots, the growth and senescence of leaves and stems, biomass accumulation, and the growth of maize grain yield as a function of soil and weather conditions, crop management practices, and variety characteristics. The model uses a standardized system for model inputs and outputs that have been described elsewhere [18 and 19]. The input system enables the user to select crop genotype (variety), weather, soil, and management data appropriate to experiment being simulated. Required crop genetic inputs for CERES Maize are given in Table 1.

**Table 1:** Definition of DSSAT maize genotype specific parameters

| **Coefficient** | **Description** |
| --- | --- |
| **P1 (°C day)** | Thermal time from seedling emergence to the end of juvenile phase |
| **P2 (days)** | Delay in development for each hour that day-length is above 12.5 hours |
| **P5 (°C day)** | Thermal time from silking to time of physiological maturity |
| **G2 (#)** | Maximum kernel number per plant |
| **G3 (mg day<sup>-1</sup>)** | Kernel growth rate during linear grain filling stage under optimum conditions |
| **PHINT (°C day tip<sup>-1</sup>)** | Thermal time between successive leaf tip appearance |

### 2.2 Field Experiments

Three sets of data were used in this study: calibration experiments, breeder evaluation experiments and model validation experiments.

The calibration experiments were conducted during the rainy and dry season of 2016 in four locations in northern Nigeria. The experiments were conducted at the Teaching and Research Farm of the Faculty of Agriculture, Bayero University, Kano (N11.516 E8.516 466m asl), at the Teaching and Research Farm of Audu Bako College of Agriculture Dambatta (N12.333 E8.517 442m asl), at the Irrigation Research Farm of Institute for Agricultural Research (IAR) Samaru, Zaria (N11.187 E7.147 702m asl) and at the Agricultural Research Station of the Kaduna Agricultural Development Project (KADP) in Saminaka, Lere (N10.52 E8.472 786 asl). Eight experiments were used for the calibration spanning over four locations, two seasons and eight planting dates (Table 2). The calibration experiment consisted of 20 varieties, but we focused on 10 varieties that were common to both the on-station calibration experiments and breeder varietal evaluation experiments (Table 3). The calibration experiments were conducted near irrigation facilities so as to maintain optimum moisture by irrigating when the soil moisture is below field capacity. Moisture conditions were monitored using a Time Domain Reflectometry (TDR) Meter 6050X1 TRASE SYSTEM (Soilmoisture Equipment Corp.). Recommended levels of mineral fertilizers for the region were applied (120N:60P:60K kg ha^−1^); potassium (K) was applied in form of Muriate of Potash, phosphorus in the form of Single Super Phosphate, and Nitrogen was applied in the form of Urea. While all the P and K fertilizers were applied at sowing; only half of the N fertilizer was applied at the time of sowing and the other half applied 21 days later. In addition, poultry manure (approximately NPK 1.1:0.8:0.5) was added to the fields at the rate of 5 tons ha^−1^ to maintain optimum nutrient status. The calibration experiments were laid down in a Randomized Complete Block Design (RCBD) with four replications. The gross plot consisted of six ridges, 0.75 m apart and 3 m long (plot area =13.5 m^2^). The two innermost ridges were used as the net plot for yield assessment and for sampling purposes. A space of 0.5 m was used between plots and 1m between replications. The experimental fields were cleared, harrowed, ridged and thereafter sprayed with a pre-emergence herbicide, Primextra (Atrazine + Metolachlor) at the rate of 4lha^−1^ before planting. The maize was sown at intra-row spacing of 0.25m at two seeds per hole, and later thinned to one plant giving a population of 53, 333 plants ha^−1^.

**Table 2:** Description of sites for field experiments and breeder evaluation data

| **Site and Environment** | **Code** | **Sowing Date** | **Ecology\*** | **Dominant Soil Type** | **Cumulative Rainfall + Irrigation (mm)** |
| --- | --- | --- | --- | --- | --- |
| **On-station Experiments for calibration** |  |  |  |  |  |
| **Bayero Uni. Farm Dry Season** | BUK DS | 16-03-2016 | SS | Typic Kandiustalf | 843 |
| **Bayero Uni. Farm Dry Season** | BUK RS | 25-07-2016 | SS | Typic Kandiustalf | 705 |
| **Dambatta Dry Season** | DBT DS | 19-03-2016 | SS | Typic Kanhaplustalf | 976 |
| **Dambatta Rainy Season** | DBT RS | 26-07-2016 | SS | Typic Kanhaplustalf | 690 |
| **Samaru Dry Season** | SMR DS | 22-03-2016 | NGS | Plinthic Haplustult | 840 |
| **Samaru Rainy Season** | SMR RS | 29-07-2016 | NGS | Plinthic Haplustult | 850 |
| **Lere Dry Season** | LER DS | 17-03-2016 | NGS | Plinthic Kandihumult | 964 |
| **Lere Rainy Season** | LER RS | 31-07-2016 | NGS | Plinthic Kandihumult | 1054 |
| **Breeder Varietal Evaluation experiments** |  |  |  |  |  |
| **Zaria 2012** | ZRA 12 | 12-06-2012 | NGS | Typic Kandiustalf | 1123 |
| **Zaria 2013** | ZRA 13 | 10-06-2013 | NGS | Typic Kandiustalf | 1222 |
| **Mokwa 2012** | MKW 12 | 08-06-2012 | SGS | Oxic Haplustult | 1346 |
| **Mokwa 2013** | MKW 13 | 28-05-2013 | SGS | Oxic Haplustult | 1402 |
| **Bagauda 2012** | BGD 12 | 13-06-2012 | SS | Typic Kandiustalf | 882 |
| **Bagauda 2013** | BGD 13 | 21-06-2013 | SS | Typic Kandiustalf | 941 |
| **Batsari 2012** | BTR 12 | 22-06-2012 | SS | Ustoxic Dystropept | 806 |
| **Batsari 2013** | BTR 13 | 21-06-2013 | SS | Ustoxic Dystropept | 854 |
| **Samaru 2012** | SMR 12 | 11-06-2012 | NGS | Typic Plinthiustalfs | 1118 |
| **Samaru 2013** | SMR 13 | 14-06-2013 | NGS | Typic Plinthiustalfs | 1241 |
| **Minjibir 2012** | MJB 12 | 21-06-2012 | SS | Typic Kandiustalfs | 791 |
| **Minjibir 2013** | MJB 13 | 18-06-2013 | SS | Typic Kandiustalfs | 824 |
| **Kadawa 2012** | KDW 12 | 23-06-2012 | SS | Typic Plinthiustalfs | 891 |
| **Kadawa 2013** | KDW 13 | 19-06-2013 | SS | Typic Plinthiustalfs | 913 |
| **Experiments for Model Evaluation** |  |  |  |  |  |
| **BUK 2013** | BUK 13 |  | SS | Typic Kandiustalfs | 892 |
| **BUK 2014** | BUK 14 |  | SS | Typic Kandiustalfs | 967 |
| **BUK 2015** | BUK 15 |  | SS | Typic Kandiustalfs | 1021 |
| **BUK 2016** | BUK 16 |  | SS | Typic Kandiustalfs | 972 |

\* SS = Sudan Savanna, NGS = Northern Guinea Savanna, SGS = Southern Guinea Savanna

**Table 3:** Characteristics of maize varieties used in the study

| S/N | Name | Common Name | Type | Maturity | Tolerance |
| --- | --- | --- | --- | --- | --- |
| 1 | 2011TZEWDTSRSTRYN | Early White | OPV¶ | Early | Drought/Striga |
| 2 | 2013TZEEWPOPDSTR | E.E White | OPV | Extra Early | Drought/Striga |
| 3 | EVDTS-W-99STR | Sammaz 32 | OPV | Early | Drought |
| 4 | EVDTS-Y-2000-STR | Sammaz 34 | OPV | Early | Drought/Striga |
| 5 | OBA SUPER 9 | Oba 9 | Hybrid | Late | - |
| 6 | M0926-8 | Seedco White | Hybrid | Late | MSV* |
| 7 | TZE124 x TZE125 | Sammaz 41 | Hybrid | Early | MSV |
| 8 | TZEE129 x TZEE121 | Ife hybrid 5 | Hybrid | Extra Early | Drought |
| 9 | TZEE-WPOPSTRC5 x TZEEI6 | Ife hybrid 6 | Hybrid | Extra Early | Drought/Striga |
| 10 | TZEYPOPDSTRC4 x TZEEI13 | Sammaz 42 | Hybrid | Extra Early | Drought/Striga |

¶ Open pollinated variety \* Maize Streak Virus

For calibration using data from breeder evaluation trials, we collected long-term yield evaluation data from breeders at the International Institute for Tropical Agriculture (IITA), Ibadan and the Institute for Agricultural Research (IAR), Zaria. Data for the 10 maize varieties used in this study were selected. The bulk data was subjected to various quality checks. We used data for the 2012 and 2013 seasons from seven locations where weather and soil data were available. Table 2 shows the locations and data used in the calibration with breeder data.

For model validation, field experiments were conducted at the Research Farm (11°59’N, 8°25’E 466m above sea level) of the Faculty of Agriculture, Bayero University, Kano in the rainy seasons between 2013 to 2016 (four seasons). The treatments consisted of three rates of nitrogen (0, 60 and 120 kg N ha^−1^) and ten maize varieties used in the calibration experiments (Table 2). Treatments were laid out in a split-plot design with three replications. Nitrogen rates were assigned to the main plots while the varieties were assigned to the sub-plot. Although the experiments were conducted in the rainy season, moisture contents were monitored and supplementary irrigation was provided to ensure no moisture stress. All conventional agronomic cultural practices were followed. The data collected for model evaluation includes grain yield (kg ha^−1^), total grain nitrogen (kg ha^−1^), total tissue nitrogen (kg ha^−1^) and nitrogen harvest index (percentage). Total grain and tissue nitrogen were determined using the Micro Kjeldahl method.

### 2.3 Plant measurements

Evaluation of crop development was done by observing the phenology of the different maize varieties, and recording the length of time (days) it takes to attain each phenological phase. The measurements were then converted to growing degree days (GDD) using the relationship:

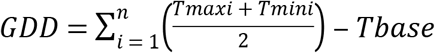

Ten plants were tagged from the center of each plot in each replication for phenological observations. The end of the juvenile stage (i.e. panicle initiation) was determined through destructive sampling and dissection of three plants, followed by observation of apical meristem to check for floral bud development at 2 d intervals starting 14d after emergence. The end of juvenile stage was recorded when the male flower primordial were visible in 50% of plants examined. Days to 50% tasseling was recorded when tassels were observed on 50% of the tagged plants. Physiological maturity observations were conducted as follows: kernels were removed from the base, middle and distal end of each sampled ear daily, starting when husks begin to show signs of drying. Days to physiological maturity was recorded when 50% of the kernels in each tagged ear had formed a black layer, indicating physiological maturity.

Plant biomass was taken at four different stages: vegetative, anthesis, grain filling and physiological maturity. Five plants within a one-meter strip in a row were cut at the ground level as suggested by Ogoshi et al. [10]. Leaves were separated from the stem, chopped and dried in the shade for three days. Both stems and leaves were oven dried at 70°C for 36 - 48 hours until the sample had attained constant weight. Yield and yield component measurements were taken at harvest maturity. Plant height was measured from five randomly tagged plants within the net plot using a standard field meter rule. Other variables measured included: the number of seeds per unit area (seed # m^−2^), dry seed weight (g m^−2^), dry cob weight (g m^−2^), dry husks weight (g m^−2^), grain yield (kg ha^−1^) and stover weight at harvest (kg ha^−1^). All yield and yield component measurements were done using procedures and formulae described by Ogoshi et al. [10]. Total grain and tissue nitrogen (measured for the evaluation experiments only) were determined using the Micro – Kjeldahl method.

### 2.4 Soil and weather data

Detailed soil studies were conducted for each experimental location before planting. Soil pits were dug in each location, and soil samples were taken from each layer. The collected samples were then analyzed for pH, texture, moisture, bulk density, exchangeable potassium (K), organic matter, phosphorus (P), total nitrogen and CEC. For the detailed calibration and validation experiments, daily weather data were collected from weather stations (Watchdog 2000 Series, Spectrum Technologies) adjacent to all experimental sites. All weather stations were less than 5 km away from the experimental sites.

### 2.5 Initialization of soil and weather parameters

Daily records of minimum and maximum temperature, total solar radiation, and total rainfall are required for the CERES-Maize model weather initialization. The *Weatherman* utility in DSSAT was used to input the weather data to create the weather file used by the CERES- Maize model. The *Weatherman* utility also requires information on name of weather station, latitude, longitude and altitude. Soil data tool (*SBuild*) was used to create the soil database which was used for the general simulation purposes. Name of the country, name of experimental site, site code, site coordinates, soil series and classification were among the data entered in this utility. Initial soil water was set to field capacity for all locations for the calibration experiments, while for the breeder evaluation this condition was not set, leaving the inputed moisture properties of the soils in each location. Measured soil characteristics taken from each profile were used to calculate the soil physical and chemical parameters that are needed to run the model. For calibration experiments, we assumed that N was not limiting while for the breeder evaluation nitrogen was simulated although N stress was not recorded in any of the locations. For the evaluation experiments however, Nitrogen was simulated and application was done according to treatments. For other simulation options, initial conditions were as reported for each year and location, the Priestly-Taylor/Ritchie method was selected for simulation of evapotranspiration while the Soil Conservation Service (SCS) method was selected for simulation of infiltration. Photosynthesis was simulated using the leaf photosynthesis response curve, while hydrology and soil evaporation were simulated using the Ritchie Water Balance and Suleiman-Ritchie methods respectively. Phosphorus and Potassium were not simulated in all trials and locations.

### 2.6 Estimating genotype specific parameters

The GENCALC program of the DSSAT (Version 4.6) was used to calibrate the GSPs of the maize varieties. GENCALC is a software package that facilitates the calculation of variety coefficients for use in existing crop models including the CSM-CERES-Maize Model [20]. The CSM-CERES-Maize model has GSPs that define growth and development characteristics or traits of a maize variety (Table 3). Three parameters (P1, P2 and P5) define the life cycle development characteristics, two coefficients (G2 and G3) define growth and yield characteristics and one coefficient, PHINT, defines leaf tip appearances [21]. All the candidate genetic coefficients were selected and calculated using GENCALC.

The varieties used in the trials were representative of all the maturity groups, i.e. extra early to late maturity. The default values in DSSAT were therefore used as initial coefficients for the extra-early, early and late maturity classes. Variety coefficient values for each variety are then varied, relative to each simulated and observed measurement. The model algorithm then searches the output file and uses the difference between simulated and observed variables to decide whether to increase or decrease the value of the coefficient that is being estimated. When GENCALC finds a good fit for each observation, it averages the coefficients and calculates the root mean square error (RMSE) [22]. According to each genetic parameter, the process is repeated until the best fit is selected. An interactive procedure is used by GENCALC where the user changes the variety coefficient step to minimize the errors and speed-up the convergence of the algorithm. The search finishes when the user accepts the parameters providing the lowest RMSE for a single target trait.

The approach used for the optimization procedure (Fig 1) was similar to that used by Anothai et al. [13]. This approach has not been reported for maize, especially in Sub-Saharan Africa. For calibration of maize genotypes using this procedure, four variables connected to four out of the six coefficients were directly measured (P1, P5, G2 and PHINT), while P2 and G3 of the initial genotypes were first selected and later adjusted until a good fit is observed. The generated coefficients were then used to run sensitivity analysis, using various iterations (not less than 6000 for each coefficient) to confirm the accuracy of the sequential approach. The adjustment for each target coefficient was done while all other non-target coefficients were kept constant.

**Fig 1:**
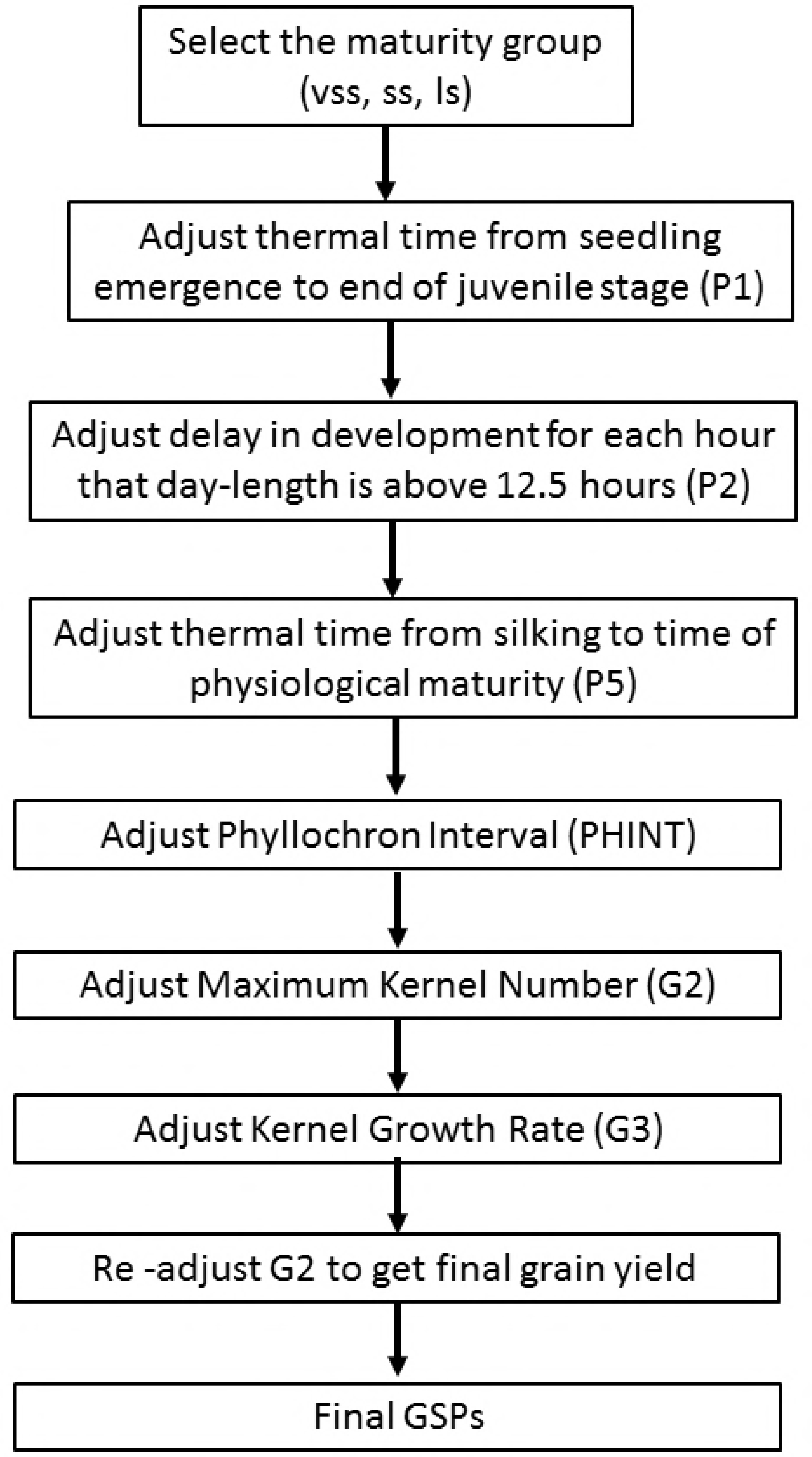
Order sequence of optimizations for calibrating the cultivar coefficients using GENCALC

### 2.7 Model evaluation

The model was calibrated using data from conventional experiments or breeder evaluation trials. Model evaluation was done using data from the nitrogen trials (Table 2). The data sets used for model evaluation were of two types; single measured data and time series data. For single measured data, we used r^2^ and root mean squared error (RMSE) (equation 1) to evaluate the agreements between simulated and observed values. Normalized Root Mean Squared Error (nRMSE, equation 2) and the index of agreement (d, equation 3) [23] were used to evaluate the time series data. We used nRMSE for time series data because RMSE varies with growth over time as the magnitude of the growth variables increase. The d-statistic was used because it gives a single index of model performance, which covers bias and variability; it also indicates 1:1 prediction better than R^2^. A low value for nRMSE (expressed in percent) is desired to define a good fit. The d statistic has values between zero and one, with one being the best fit. The modeling efficiency, EF [24] was employed to test modeling efficiency (equation 4). EF has no dimension and an EF = 1 corresponds to a perfect match between observed and simulated data. When EF < 0, the simulated values are worse than simply using the observed mean. R^2^, RMSE, RMSEn, d-index and EF are shown in equations 1, 2, 3 and 4 respectively.

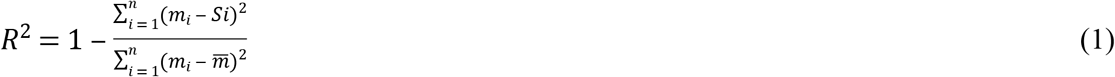

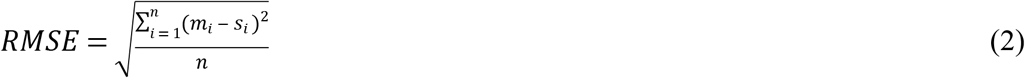

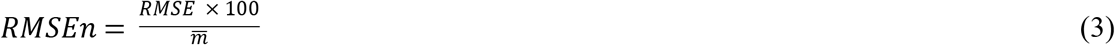

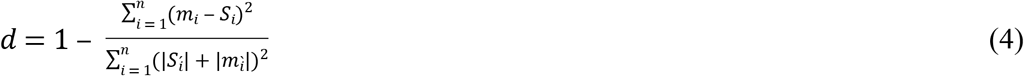

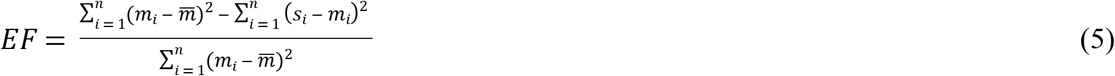

Where *n* is the number of observations, *S*_*i*_ is the simulated data, *m*_*i*_ is the measured data, and 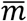 is the mean of the measured data.

## 3.0 Results

### 3.1 Calibration and breeder evaluation experiments

#### 3.1.1 Genotype specific parameters

The values of GSPs generated using data from calibration experiments and breeder evaluation data are shown in Table 4. The highest degree days from emergence to end of Juvenile stage (P1), and from silking to end of physiological maturity (P5), were recorded for SC 8325 in both the experimental and breeder data. The lowest degree days was recorded for Ife hybrid 5 using calibration data and Ife hybrid 6 for breeder data. The variety SC 8325 produced the largest number of maximum possible kernels (G2) for both data sets. The value of G3 (kernel filling rate) ranged between 6.32 and 8.20 for the experimental data, and between 6.50 and 8.40 for the breeder data. Phyllochron interval (PHINT) values ranged from 36.9 and 45.5 °Cd for the experimental data and between 34.9 and 55.0° Cd for the breeder data.

**Table 4:** Generated genotype specific parameters (GSPs) using experimental and breeder data

| Variety | P1<br>(°C day) |  | P2<br>(days) |  | P5<br>(°C day) |  | G2<br>(#) |  | G3<br>(mg day<sup>-1</sup>) |  | PHINT<br>(°C day tip<sup>-1</sup>) |  |
| --- | --- | --- | --- | --- | --- | --- | --- | --- | --- | --- | --- | --- |
|  | Expt. | Breeder | Expt. | Breeder | Expt. | Breeder | Expt. | Breeder | Expt. | Breeder | Expt. | Breeder |
| **Ife hybrid 6** | 223.6 | 269.7 | 0.500 | 0.500 | 520.7 | 520.0 | 706.7 | 519.0 | 6.93 | 6.80 | 36.90 | 34.90 |
| **Sammaz 41** | 233.6 | 280.3 | 0.500 | 0.500 | 550.7 | 530.0 | 806.9 | 766.9 | 7.80 | 7.73 | 37.00 | 42.73 |
| **Ife hybrid 5** | 213.7 | 267.8 | 0.500 | 0.500 | 511.6 | 521.4 | 518.7 | 523.6 | 7.18 | 7.20 | 40.00 | 36.30 |
| **Sammaz 42** | 230.0 | 287.6 | 0.500 | 0.500 | 683.4 | 681.7 | 786.7 | 793.6 | 7.20 | 7.86 | 45.50 | 37.80 |
| **OBA SUPER I** | 293.1 | 290.3 | 0.500 | 0.500 | 768.1 | 780.0 | 828.7 | 840.0 | 7.30 | 7.80 | 45.00 | 45.00 |
| **SC 8325** | 289.8 | 291.3 | 0.500 | 0.500 | 781.8 | 783.6 | 834.1 | 841.7 | 8.20 | 8.40 | 41.20 | 45.20 |
| **Sammaz 34** | 287.0 | 287.0 | 0.500 | 0.500 | 596.0 | 596.0 | 827.0 | 827.0 | 6.52 | 6.52 | 40.00 | 40.00 |
| **Sammaz 32** | 282.0 | 235.0 | 0.500 | 0.500 | 601.0 | 800.0 | 822.0 | 722.0 | 6.50 | 6.50 | 45.04 | 39.00 |
| **IITA Early White** | 270.0 | 190.0 | 0.500 | 0.500 | 614.3 | 660.0 | 713.4 | 780.0 | 6.40 | 8.00 | 45.00 | 55.00 |
| **IITA Extra Early White** | 183.6 | 207.6 | 0.500 | 0.500 | 601.0 | 652.3 | 523.3 | 601.4 | 6.32 | 7.12 | 42.10 | 45.00 |

#### 3.1.2 Phenology and growth

Evaluation of CERES-Maize for grain yield, number of days to anthesis, number of days to physiological maturity and plant height and using both calibration experiments and breeder evaluation is shown in Fig 2 for two varieties. Calibration of number of days to anthesis, and plant height, were more accurate when experimental data were used compared with breeder data for both varieties. Calibration of both variables using experimental data resulted in d-index values in the range of 0.83-0.96 for the trial data. For the breeder data however, d-index values ranged from 0.43 to 0.69. Days to anthesis was calibrated with higher accuracy than plant height for all varieties. Number of leaves per plant and plant height were measured for the experimental data at different time intervals. The simulated values for both plant height and number of leaves were accurate at all sampling periods (Fig 3).

**Fig 2:**
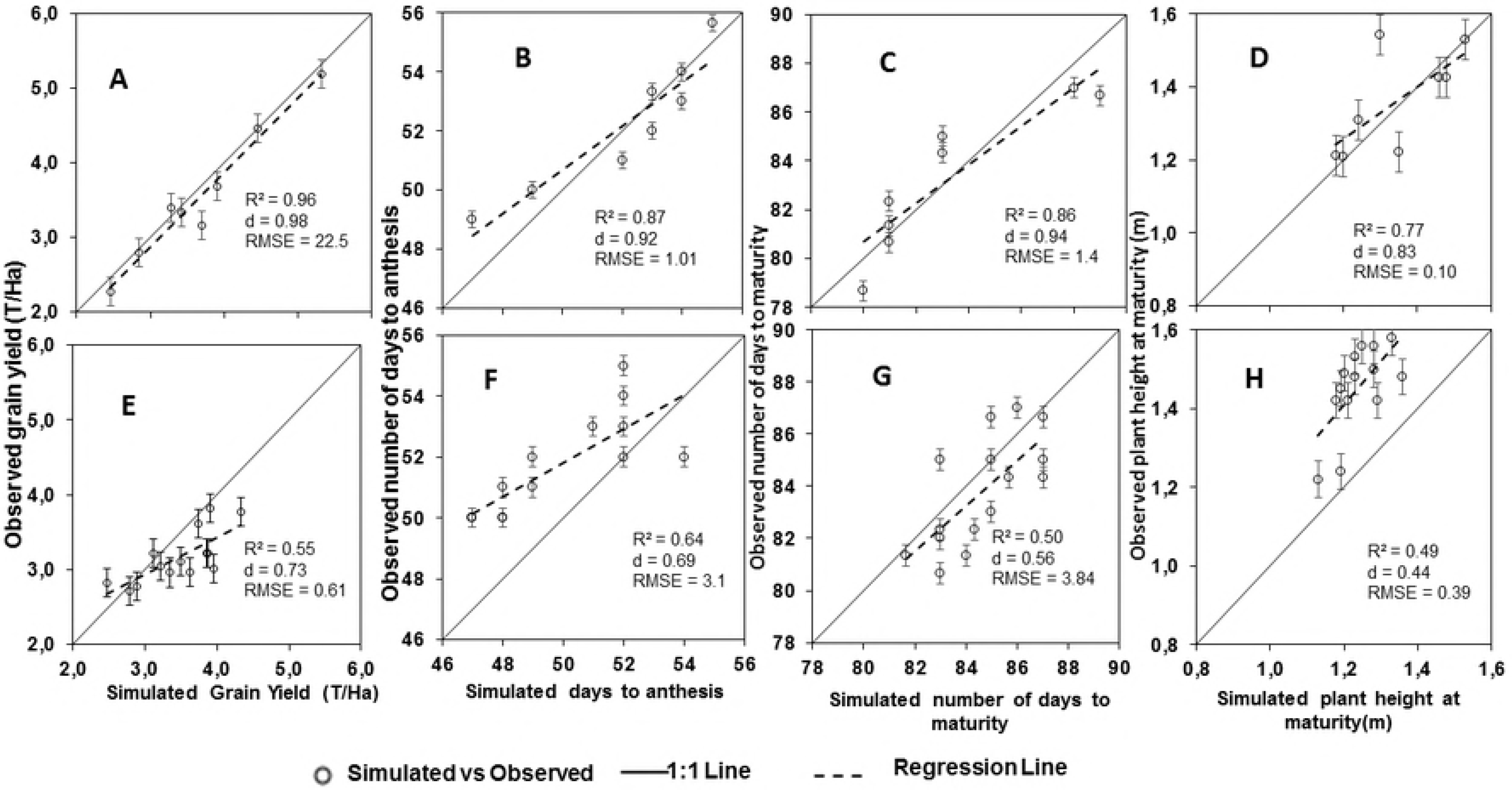
Comparisons between simulated and observed grain yield, days to anthesis, days to maturity and plant height at harvest for SAMMAZ 32 using experiment (A, B, C, D) and breeder (E, F, G, H) data. Solid lines = 1:1 lines; dashed lines = regression lines. Error bars denote Standard Error of Mean (SEM)

**Fig 3:**
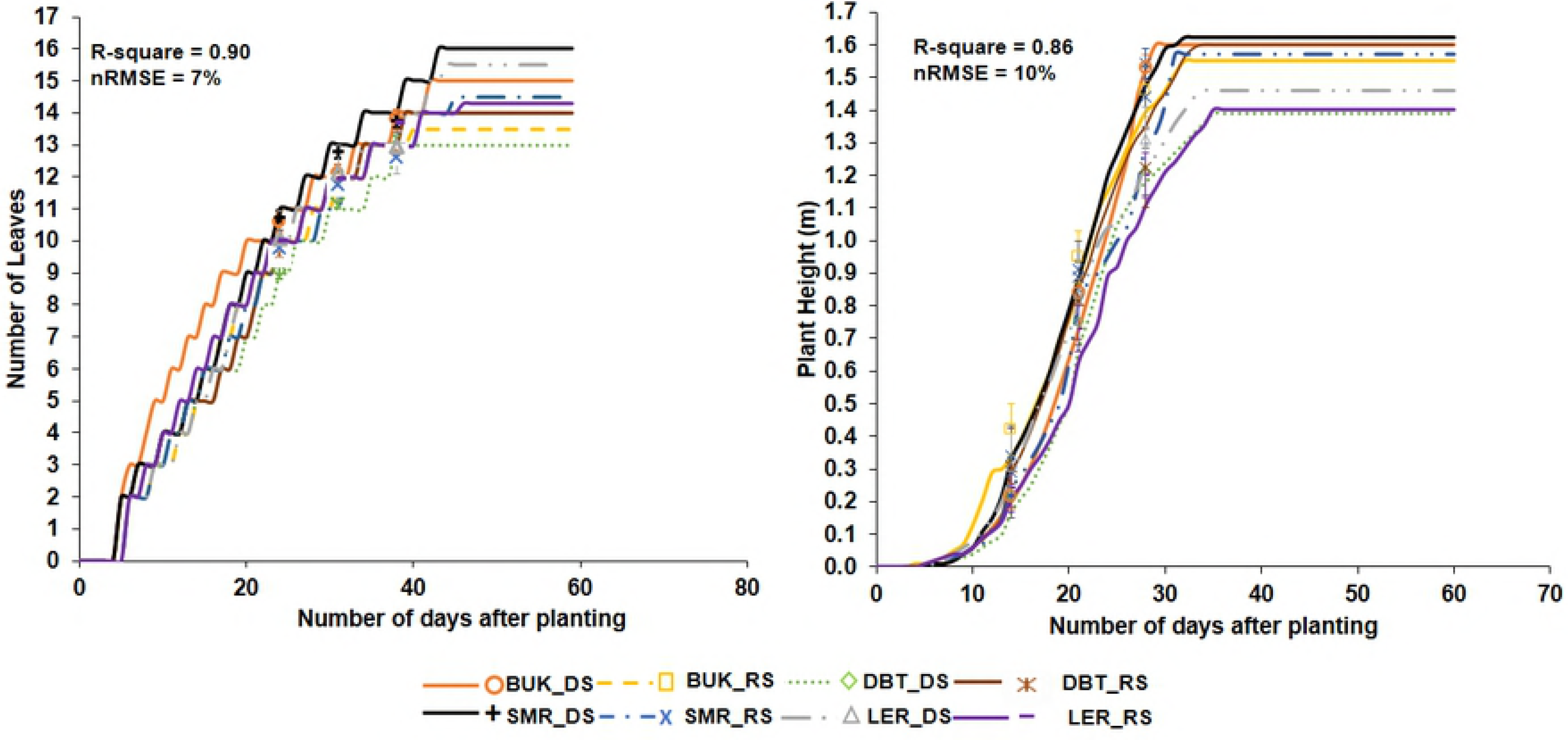
Simulated (lines) vs Observed (symbols) plant heights and number of leaves of SAMMAZ 32 using experiment data. Error bars denote Standard Error of Mean (SEM)

#### 3.1.3 Biomass and Leaf Area Index

Biomass and LAI were measured at juvenile stage, at anthesis, and at physiological maturity for the calibration data only. Fig 4 shows the result of simulation of above-ground biomass and LAI for Sammaz 32 across the trial locations. Good agreements were found between simulated and observed variables for all other varieties. Biomass was simulated with higher accuracy than LAI across all locations. Simulation of both biomass and LAI were most accurate using data from Samaru (d-index = 0.94, RMSE = 583.9 for biomass and d-index 0.93, RMSE 0.02 for LAI). Calibration of both variables had the lowest accuracy at Dambatta. Agreements between observed and simulated LAI were closer for the earliest measurement (juvenile stage), followed by measurement at anthesis, and physiological maturity in all locations except at Samaru where the reverse was observed. For biomass however, measurement at physiological maturity produced the closest agreements between observed and simulated values, while measurement at anthesis produced the lowest agreement between observed and simulated variables.

**Fig 4:**
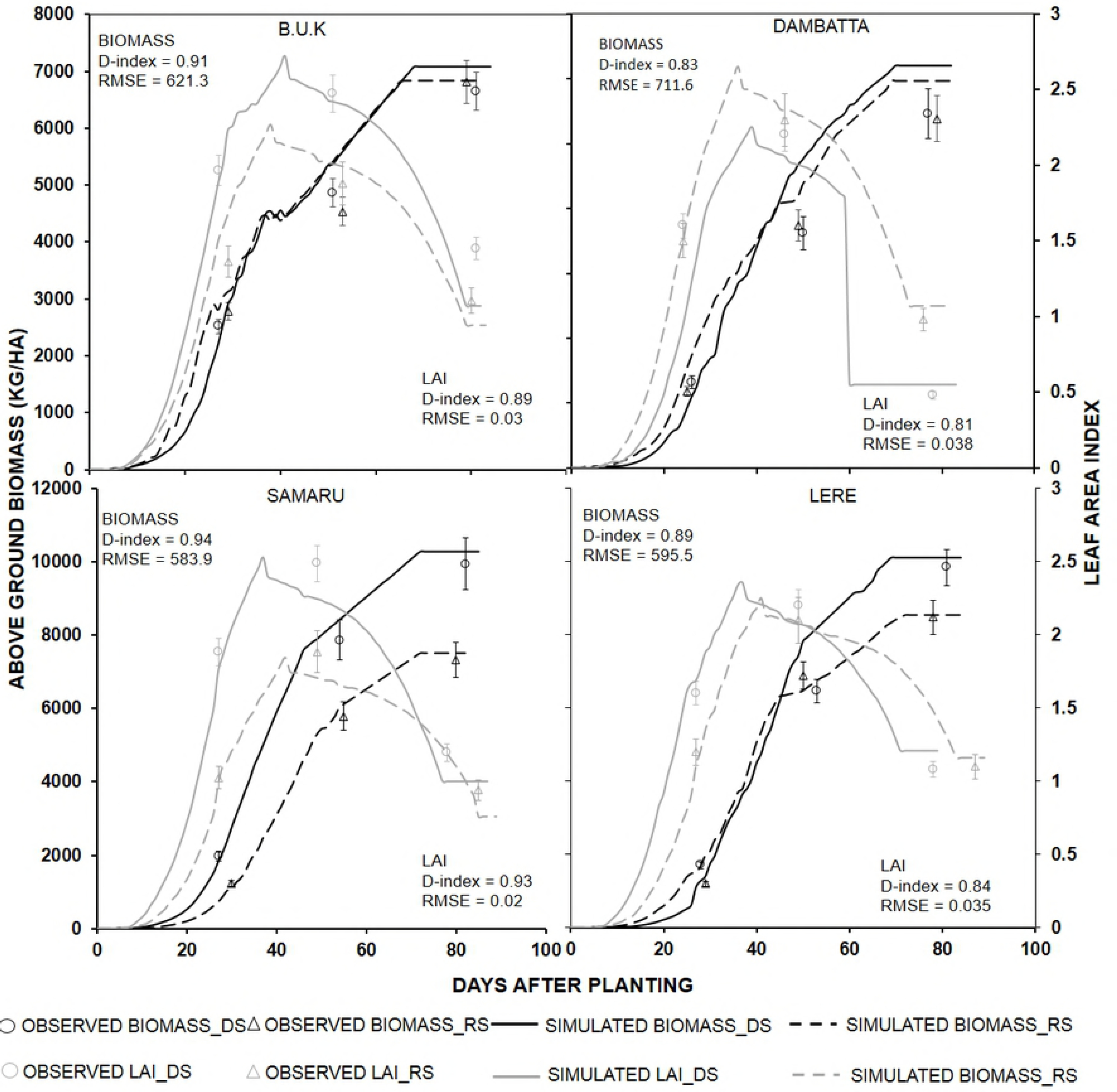
Simulated (lines) vs Observed (figures) Biomass and LAI of SAMMAZ 32 using experimental data. Error bars denote Standard Error of Mean (SEM)

#### 3.1.4 Yield and yield attributes

Yield and yield attributes were well calibrated for all varieties in both calibration and breeder datasets. Table 5 shows the result of comparisons between observed and simulated mean grain yields of all varieties across different locations. Calibration of grain yield using experimental data was more accurate, as evidenced by low percentage prediction deviations (1.2 to 14.1). Values for model statistics were also good for the calibration data (RMSE = 259 kg ha^−1^, nRMSE = 6.5%, and d-index = 0.98). For the breeder data however, prediction deviations of up to 23.7% were observed, with higher RMSE (609 kg ha^−1^) and nRMSE (19.3%).

**Table 5:** Observed and simulated mean grain yields (kg ha^−1^) of all varieties across different locations

| Data Type | Observed* | Simulated | PD%# |
| --- | --- | --- | --- |
| **Experiment Data** |  |  |  |
| **BUK_DS** | **3828** | **4260** | **10.1** |
| **BUK_RS** | **3209** | **3640** | **11.8** |
| **DBT_DS** | **2758** | **3054** | **9.7** |
| **DBT_RS** | **2628** | **2883** | **8.9** |
| **SMR_DS** | **5030** | **5476** | **8.1** |
| **SMR_RS** | **3536** | **3939** | **10.2** |
| **LERE_DS** | **4561** | **4863** | **6.2** |
| **LERE_RS** | **3452** | **4095** | **15.7** |
| **RMSE (kg/ha)** | **416.6** |  |  |
| **nRMSE (%)** | **11.4** |  |  |
| **EF** | **0.89** |  |  |
| **d-index** | **0.94** |  |  |
| **Breeder Data** |  |  |  |
| **ZRA 12** | **2958** | **3345** | **11.6** |
| **ZRA 13** | **2969** | **3625** | **18.1** |
| **MKW 12** | **3214** | **3866** | **16.9** |
| **MKW 13** | **3042** | **3213** | **5.3** |
| **BGD 12** | **3812** | **3913** | **2.6** |
| **BGD 13** | **2782** | **2885** | **3.6** |
| **BTR 12** | **3226** | **3110** | **-3.7** |
| **BTR 13** | **3112** | **3487** | **10.8** |
| **SMR 12** | **3214** | **3863** | **16.8** |
| **SMR 13** | **3779** | **4329** | **12.7** |
| **MJB 12** | **3612** | **3746** | **3.6** |
| **MJB 13** | **2831** | **2470** | **-14.6** |
| **KDW 12** | **2711** | **2779** | **2.4** |
| **KDW 13** | **3017** | **3956** | **23.7** |
| **RMSE (kg/ha)** | **609.1** |  |  |
| **nRMSE (%)** | **19.3** |  |  |
| **EF** | **-0.45** |  |  |
| **d-index** | **0.73** |  |  |

\* Mean for all varieties; # Percentage prediction deviation

### 3.2 Model validation experiments

Grain and tissue nitrogen, as well as grain yield, at harvest were simulated using independent datasets from trials conducted at BUK during the rainy seasons between 2013 and 2016. Simulations were done using GSPs generated from both experimental and breeder data. Table 6 shows the comparison between observed and simulated grain yields with accompanying model statistics for the two datasets taking SAMMAZ 32 and EE-White as examples. Grain yield was well simulated for both varieties using both datasets, although better fits were observed for GSPs from the calibration data. Nonetheless, low values of RMSE (below 3% of mean for experimental and 5% for breeder), high values of d index (0.99 for experimental and 0.96 for breeder) and good EF values (slightly less thqn 1 for both datasets) were observed. Tables 7 and 8 shows comparisons of simulated grain and stover nitrogen using GSPs generated from calibration and breeder evaluation experiments. Better agreements between observed and simulated grain and stover Nitrogen were observed at high Nitrogen (120 and 60 Kg N) for both calibration and breeder evaluation experiments. At zero nitrogen application however, the agreements between observed and simulated values where low as evidenced by higher RMSE and lower d-index values.

**Table 6:** Simulated vs Observed grain yields of Sammaz 32 and EE White in the model validation experiments, under different nitrogen levels using GSPs derived from calibration experiment and breeder evaluation experiment

| Treatment | Observed | Simulated<br>(GSPs < Calibration<br>experiment) | Simulated<br>(GSPs < Breeder<br>evaluation trials) |
| --- | --- | --- | --- |
| **Sammaz 32** |  |  |  |
| **0 kg N** | 1245 | 1318 | 1069 |
| **60 kg N** | 2648 | 2635 | 2665 |
| **120 kg N** | 3255 | 3142 | 2817 |
| ***SE*±** | 57.3 |  |  |
| ***RMSE*** |  | 47.8 | 272.7 |
| ***D-Index*** |  | 0.99 | 0.97 |
| ***EF*** |  | 0.91 | 0.89 |
| **EE White** |  |  |  |
| **0 kg N** | 979 | 962 | 1174 |
| **60 kg N** | 2177 | 2015 | 2512 |
| **120 kg N** | 3092 | 2996 | 3377 |
| ***SE*±** | 60.6 |  |  |
| ***RMSE*** |  | 109.2 | 277.7 |
| ***D-Index*** |  | 0.99 | 0.97 |
| ***EF*** |  | 0.98 | 0.89 |

**Table 7:** Comparison of simulated and observed grain nitrogen (kg N ha^−1^) of SAMMAZ 32 for GSPs generated using calibration experiments and breeder evaluation experiments

|  | **SIM (Calibration Experiments)** | **SIM (Breeder Evaluation Expts.)** | **OBS** |
| --- | --- | --- | --- |
| **120 kg N ha<sup>-1</sup>** |  |  |  |
| **BUK 13** | 43.0 | 45.3 | 42.1 |
| **BUK 14** | 41.5 | 48.3 | 44.3 |
| **BUK 15** | 45.1 | 46.4 | 42.2 |
| **BUK 16** | 41.1 | 44.9 | 43.3 |
| ***SE*±** |  |  | 0.81 |
| ***RMSE*** | 1.46 | 3.18 |  |
| ***d-index*** | 0.65 | 0.41 |  |
| **60 kg N ha<sup>-1</sup>** |  |  |  |
| **BUK 13** | 42.8 | 45.0 | 43.2 |
| **BUK 14** | 45.1 | 46.2 | 43.3 |
| **BUK 15** | 40.1 | 44.4 | 41.0 |
| **BUK 16** | 45.5 | 43.0 | 43.6 |
| ***SE*±** |  |  | 0.79 |
| ***RMSE*** | 1.17 | 2.8 |  |
| ***d-index*** | 0.77 | 0.49 |  |
| **0 kg N ha<sup>-1</sup>** |  |  |  |
| **BUK 13** | 12.1 | 9.4 | 14.3 |
| **BUK 14** | 25.1 | 24.1 | 21.8 |
| **BUK 15** | 26.8 | 30.6 | 20.6 |
| **BUK 16** | 6.9 | 5.5 | 10.2 |
| ***SE*±** |  |  | 0.94 |
| ***RMSE*** | 4.6 | 5.1 |  |
| ***d-index*** | 0.68 | 0.62 |  |

**Table 8:** Comparison of simulated and observed stover nitrogen (kg N ha^−1^) of SAMMAZ 32 for GSPs generated using data from calibration experiments and breeder evaluation experiments

|  | **SIM (Calibration Experiments)** | **SIM (Breeder Evaluation Expts.)** | **OBS** |
| --- | --- | --- | --- |
| **120 kg N ha<sup>-1</sup>** |  |  |  |
| **BUK 13** | 80.1 | 126.8 | 78.7 |
| **BUK 14** | 78.8 | 135.9 | 76.5 |
| **BUK 15** | 77.4 | 130.8 | 72.6 |
| **BUK 16** | 78.4 | 112.9 | 76.3 |
| ***SE*±** |  |  | 17.8 |
| *RMSE* | 3.48 | 12.8 |  |
| *d-index* | 0.83 | 0.21 |  |
| **60 kg N ha<sup>-1</sup>** |  |  |  |
| **BUK 13** | 64.9 | 75.2 | 67.8 |
| **BUK 14** | 81.4 | 95.0 | 77.4 |
| **BUK 15** | 86.7 | 91.9 | 78.8 |
| **BUK 16** | 70.8 | 64.4 | 62.6 |
| *SE*± |  |  | 4.6 |
| *RMSE* | 6.4 | 8.2 |  |
| *d-index* | 0.68 | 0.71 |  |
| **0 kg N ha<sup>-1</sup>** |  |  |  |
| **BUK 13** | 27.4 | 25.1 | 23.2 |
| **BUK 14** | 30.2 | 45.7 | 27.4 |
| **BUK 15** | 40.2 | 49.8 | 30.6 |
| **BUK 16** | 16.9 | 18.2 | 14.3 |
| *SE*± |  |  | 2.47 |
| *RMSE* | 7.16 | 15.5 |  |
| *d-index* | 0.63 | 0.65 |  |

## 4.0 Discussion

Calibrated GSPs from the on-station experiments and breeder evaluation experiments were similar to GSPs reported for related varieties in West and Southern Africa [29–31] with respect to yield and yield attributes. For growth and phenology however, data from our experiments produced better calibration of growth and phenology than earlier reported experiments in the Nigerian Savannas. Calibration of the GSPs using the on-station experiment datasets produced better model fits than the breeder evaluation data as expected. The closeness of fit observed for the on-station data could be attributed to better experimental sites (soils with higher fertility and better moisture retention), better crop management (timely weeding, fertilizer application etc.) and higher experimental precision. This is evidenced by the breeder data having higher experimental errors for all measured variables when compared to the evaluation experiments. The evaluation experiments were also done on larger plot sizes and no missing plants were recorded at harvest, while in the breeder data smaller plots were used and there was no considerations for missing plants during yield calculations. In addition, for the experimental datasets more plant-related variables were measured compared to the breeder evaluation experiment data where only grain yield, days to flowering, plant height and days to physiological maturity were measured. For the breeder evaluation experiment, the closeness between observed and simulated plant heights was low. This could be attributed to the fact that most breeder trials are conducted under water limited conditions, thus rainfall variability may affect crop performance and data quality. Although the model is able to properly simulate water stress, no stress was observed in any of the breeder evaluation sites and years. Grain yield and days to anthesis were simulated more accurately than plant height for the breeder evaluation experiment. This can be attributed to the high number of datasets used (7 locations and 2 seasons). Anothai et al. [32] suggested that more accurate predictions of yield and phenology are observed when data is collected from many locations and seasons. For the on-station experiment, plant height, number of leaves, leaf area index, biomass, number of grains per meter square and grain yields were well calibrated as the differences between observed and simulated values were very minimal.

According to literature (32, 33, and 35) when many years and locations are available, GSPs calibrated using breeder evaluation experiments produced very accurate comparisons between observed and simulated growth, yield and phenology of maize. As suggested by Fensterseifer et al. [33], uncertainties exist in the reliability of model based simulations of growth, yield and phenology when calibrations are done using data from trials conducted under few environmental conditions. Also, several researchers [34–35] reported that the major factors determining the success of a model calibration process, which determines the applicability of the model on a larger scale is dependent on the wide variability of data used during the calibration process. Thorp et al. [36] suggested that for accurate calibration of crop models, integration of time variation using different planting dates and seasons, and spatial differences using different locations of datasets should be adopted for calibration of crop models using datasets from yield/breeder evaluation trials. To further verify these claims, we re-ran a couple of contrasting varieties under both on-station experiments and breeder evaluation experiments using different number of trials and data sets. For the on-station experiments, we first of all reduced the number of experiments by subtracting 2 stations concurrently (i.e. reducing from 8-6, 6-4 and 4-2). With every decrease in number of experiments, a subsequent decrease in model efficiency and increase in prediction error were recorded. The higher the number of trials the better the model fitted the observations, also reducing the number of experiments to 4 led to EF and d-index values below 0.4, while further reduction to 2 reduced the model efficiency to 0.25 and increased the prediction error to 55%. Using 4 experiments and all measured data produced the lowest level of acceptable model statistics (d-index ≥ 0.45, nRMSE ≤ 15% and EF ≥0.4). For the breeder evaluation experiments, every reduction in number of experiments led to a decrease in model efficiency and an increase in prediction error. We also reduced the number of datasets from the evaluation experiments to exactly the same that was used in the breeder evaluation experiments. This resulted in decrease in model efficiency (0.87 to 0.79 for Sammaz 32 and from 0.92 to 0.87 for Seedco white). This shows that the number of experimental sites are more important than amount of calibration data as long as the minimum data sets (MDS) are collected as shown in Table 9. This view is supported by [33 and 29]. When a large number of locations and planting dates are available, data from breeder evaluation experiments in the SSA can be used to make good calibration of calibrations with lower RMSE & nRMSE and higher d-index & EF values.

**Table 9:**
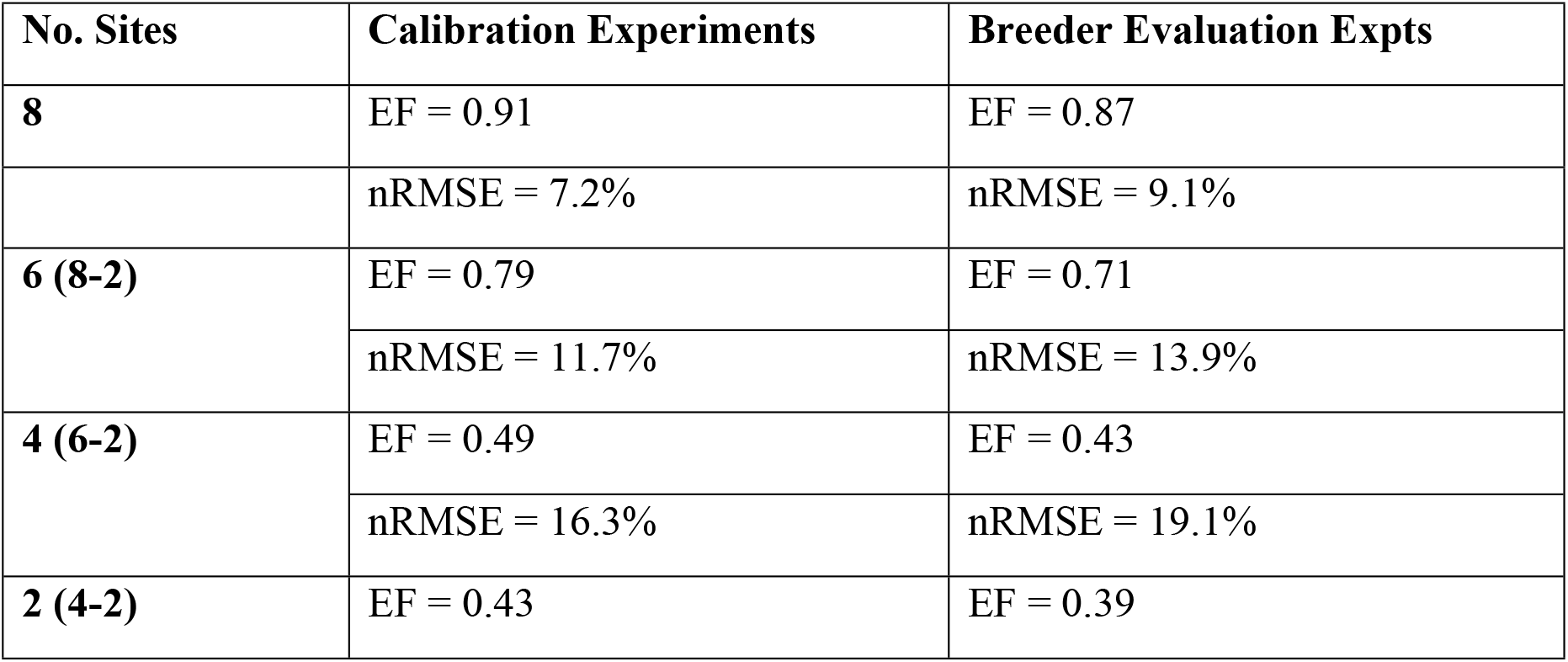
Model statistics values for reduction in number of experimental sites for both calibration experiments and breeder evaluation experiments

Although the calibration experiments provided more accurate GSPs, they are still very expensive and laborious and thus are nearly impossible to carry out especially in Sub-Saharan Africa where expertise and resources are limiting. Breeder evaluation data could also be used for calibration of GSPs where such data is available. As shown earlier, very accurate GSPs could be generated if large amount of data from many years (also planting dates) and various locations are available. This will go a long way in providing model users with cheap and easy ways of calibrating GSPs of existing and newly released varieties to their locations.

Evaluating the generated GSPs for simulation of grain yield, tissue nitrogen and grain nitrogen using independent datasets resulted in good agreements between observed and simulated values. For grain yield, comparisons of measured and simulated values using both GSPs generated using experimental and breeder data showed very close agreements under medium and high nitrogen applications. For comparisons under nitrogen stressed conditions however, poor agreements existed between observed and simulated grain yields for both GSPs. This is a common occurrence with simulations of grain yield and yield attributes under low nitrogen fertilizer applications. Gungula et al. [27], reported that the CERES-Maize model poorly predicts performance of maize under low nitrogen conditions in the tropics. The agreements between observed and simulated grain and stover nitrogen for both GSPs under high fertilizer applications is an indication that CERES-model still performs best under high nitrogen applications especially on tropical soils.

The CERES-Maize model has been shown over the years to be an important tool in evaluating crop management [3], climate change impacts [26], fertilizer recommendations [27 and 6] and yield forecasting [28]. Calibrating the newly released maize varieties currently recommended for the Nigerian maize belts will provide an important input requirement for using crop models to evaluate major production constraints including optimum stand density (OSD), appropriate varietal selection (targeting/stability analysis), choice of major partner crop (in case of mixed cropping) and fertilizer (especially N and P) managements. The availability of accurate GSPs for all major varieties will also increase the applicability of the model on a wider scale and for broader applications.

## 5.0 Conclusion

Financial as well as time constraints coupled with frequent release of new varieties makes it difficult for model users to conveniently calibrate GSPs of crop models using detailed calibration experiments. Large numbers of evaluation trials are conducted across multiple locations under diverse planting dates by breeders and other growers prior to varietal release. Availability of such datasets, especially from evaluation trials conducted under minimal stress (moisture and nutrient) conditions provides an opportunity for efficient and rapid means of generating GSPs of newly released maize varieties. A systematic approach (as proposed in this study) as well as availability of large datasets from different locations and planting dates provide opportunities for estimation of accurate GSPs. Although it is possible to generate GSPs from breeder evaluation data, care must be taken to collect data from trials conducted under optimal conditions and not too far away from a weather stations. Also, breeder data to be used for calibration of crop models must be collected from sites where detailed soil data is available. Availability of GSPs of new varieties as soon as they are released will help farmers and growers to make improved site-specific decision support tools (DST). Also, researchers will be provided with new ways to making variety groupings as well as studying complex Genotype, Environment, Management (GxExM) interactions. Model users should endeavor to join breeding units/teams in order to ensure collection of robust data needed for model calibrations that are not traditionally collected by breeders.

## 6.0 Acknowledgement

The work was supported with funding from the International Maize and Wheat Improvement Centre (CIMMYT) under the Taking Maize Agronomy to Scale in Africa (TAMASA) project. TAMASA is made possible by the generous support from Bill and Melinda Gates Foundation (contract OPP1113374). Any opinions, findings, conclusion, or recommendations expressed in this publication are those of the authors and do not necessarily reflect the view of the donor.

## Reference

1. Badu-Apraku, B., Fakorede M. A. B., Lum A. F., and Akinwale R. Improvement of yield and other traits of extra-early maize under stress and non-stress environments. Agron Journ. 2009; 101: 381–389.

2. FAO 2016, Production Year Book. Food and Agriculture Organization of the United Nations, Rome, Italy.

3. MacCarthy DS, Adiku SGK, Freduah BS, Gbefo F and Kamara AY. Using CERES-Maize and ENSO as Decision Support Tools to Evaluate Climate-Sensitive Farm Management Practices for Maize Production in the Northern Regions of Ghana. Front Plant Sci. 2017; 8:31. doi: 10.3389/fpls.2017.00031

4. Zinyengere N, Crespo O, Hachigonta S, Tadross M. Crop model usefulness in drylands of southern Africa: an application of DSSAT. S Afr J Plant S. 2015; 32(2): 95–104. Doi:10.1080/02571862.2015.1006271

5. Adnan AA, Jibrin JM, Kamara AY, Abdulrahman BL, Shaibu AS and Garba II. CERES-Maize Model for Determining the Optimum Planting Dates of Early Maturing Maize Varieties in Northern Nigeria. Front Plant Sci. 2017; 8:1118. doi:10.3389/fpls.2017.01118

6. Adnan AA, Jibrin JM, Kamara AY, Abdulrahman BL, Shaibu AS. Using CERES-Maize model to determine the nitrogen fertilization requirements of early maturing maize in the Sudan Savanna of Nigeria, J Plant Nutr. 2017;40:7, 1066–1082, DOI:10.1080/01904167.2016.1263330

7. IBSNAT. Decision Support System for Agrotechnology Transfer (DSSAT). User guide. Version 3. University of Hawaii, Honolulu. HI. 1994

8. Román-Paoli E., Welch SM, and Vanderlip RL. Comparing genetic coefficient estimation methods using the CERES-Maize model. Agr Syst, 2000;65 29–41

9. Jones CA, Kiniry JR. CERES-Maize: A Simulation Model of Maize Growth and Development. Texas A&M University Press, College Station, TX. 1986.

10. Ogoshi RM, Cagauan BG, Tsuji GY. Field and Laboratory Methods for Collection of Minimum Data sets. In: Hoogenboom, G., Wilkens P.W. and Tsuji G.Y., Eds., DSSAT, Version 3, Vol. 4. IBSNAT-ICASA, University of Hawaii, Honolulu, 217–286.1999

11. Banterng, P, Patanothai A, Pannangpetch K, Jogloy S, Hoogenboom G. Determination of genetic coefficients for peanut lines for breeding applications. Eur J Agron. 2004;21: 297–310.

12. du Toit, AS. Comparisons of using fitted (calculated) and determined (measured) genetic coefficient G2 in CERES-Maize. S Afr J Plant S. 2002; 19:4, 208–210, DOI:10.1080/02571862.2002.10634467

13. Anothai J, Patanothai A, Jogloy S, Pannangpetch K, Boote, KJ Hoogenboom G. A sequential approach for determining the cultivar coefficients of peanut lines using end-of-season data of crop performance trials. Field Crop Res. 2008;108:169–178. http://dx.doi.org/10.1016/j.fcr.2008.04.012

14. Mavromatis T, Boote KJ, Jones JW, Irmak A, Shinde D, Hoogenboom G. Developing Genetic Coefficients for Crop Simulation Models with Data from Crop Performance Trials. Crop Sci. 2001;41:40–51.

15. Bannayan M and Hoogenboom, G. Using Pattern Recognition for Estimating cultivar Coefficients of a Crop Simulation Model. Field Crop Res. 2009;111:290–302. http://dx.doi.org/10.1016/j.fcr.2009.01.007

16. He J, Jones JW, Graham WD, Dukes MD. Influence of Likelihood Function Choice for Estimating Crop Model Parameters Using the Generalized Likelihood Uncertainty Estimation Method. Agr Syst. 2010; 103:256–264. http://dx.doi.org/10.1016/j.agsy.2010.01.006

17. Jones J. W., Hoogenboom G., Porter C. H., Boote K. J., Batchelor W. D., Hunt L. A., Wilkens P. W., Singh U., Gijsman A. J., Ritchie J. T. (2003). The DSSAT cropping system model. Eur J Agron. 18, 235–265.

18. Anothai J, Patanothai A, Jogloy S, Pannangpetch K, Boote, KJ Hoogenboom G. Reduction in data collection for determination of cultivar coefficients for breeding application. Agr Syst. 2008;96, 195–206.

19. Paz JO, Batchelor WD, Colvin TS, Logsdon SD, Kaspar TC, Karlen DL. Calibration of a crop growth model to predict spatial yield variability. T Am Soc Ag Eng. 1998; 41: 1527–1534.

20. Hunt LA, Pararajasingham S, Jones JW, Hoogenboom G, Imamura DT, and Ogoshi RM. GENCALC: Software to Facilitate the Use of Crop Models for Analyzing Field Experiments. Agron J. 1993; 85:1090–1094.

21. Boote K.J, Jones JW, Batchelor WD, Nafziger ED, Myers O. Genetic coefficients in the CROPGRO-Soybean model: links to field performance and genomics. Agron J., 2003; 95:32–51.

22. Wallach D, Goffinet B. Mean squared error of prediction in models for studying ecological and agronomic systems. Biometrics. 1987; 43: 561–573.

23. Willmott CJ. Some comment on the evaluation of model performance. B Am Meteorol Soc. 1982; 63, 1309–1313.

24. Loague K, Green RE. Statistical and graphical methods for evaluating solute transport models: overview and application. J Contam Hydrol. 1991;7, 51–73.

25. Nash, J. E.; Sutcliffe, J. V. River flow forecasting through conceptual models part I — A discussion of principles. J Hydrol. 1970; 10(3): 282–290. doi:10.1016/0022-1694(70)90255-6.

26. Reimund AC, Rötter, Reiner L, Enders A, Fronzek, S, Frank E.. Implication of crop model calibration strategies for assessing regional impacts of climate change in Europe. Agric. Forest Met. 2012;32–46.

27. Gungula DT, Kling JG, and Togun AO. CERES-maize predictions of maize phenology under nitrogen-stressed conditions in Nigeria. Agron J. 2003; 95:892–899. doi:10.2134/agronj2003.0892

28. Soler CMT, Sentelhas PC, Hoogenboom G. Application of the CSM-CERES-Maize model for planting date evaluation and yield forecasting for maize grown off-season in a subtropical environment. Eur J Agron. 2007;27: 165–177

29. Jagtap SS, Mornu M, Kang BT. Simulation of growth, development, and yield of maize in the transition zone of Nigeria. Agr Syst. 1993;41, 215–229.

30. Jibrin MJ, Kamara AY, Friday E. Simulating planting date and cultivar effect on dryland maize production using CERES maize model. Afr J Agr Res. 2012; 7: 5530–5536. doi: 10.5897/AJAR12.1303

31. Chisanga CB, Phiri E, Shepande C, Sichingabula H. Evaluating CERES-Maize Model Using Planting Dates and Nitrogen Fertilizer in Zambia. J Agric Sci, 2015;7(3);

32. Anothai J, Patanothai A, Jogloy S, Pannangpetch K, Boote, KJ Hoogenboom G. Reduction in data collection for determination of cultivar coefficients for breeding application. Agr Syst. 2008;96, 195–206.

33. Fensterseifer CA, Streck NA, Baigorria GA, Timilsina AP, Zanon AJ, Cera JC, Rocha TSM. On the number of experiments required to calibrate a cultivar in a crop model: The case of CROPGRO-soybean. Field Crop Res. 2017; 204146–152. http://dx.doi.org/10.1016/j.fcr.2017.01.007

34. Ruíz-Nogueira B, Boote KJ, Sau F. Calibration and use of CROPGRO-soybean model for improving management under rainfed conditions. Agr Syst. 2001; 68: 151–173.

35. Xiong W, Balkovic J, van der Velde M, Zhang X, Izaurralde RC, Skalsky R, Lin E, Mueller NO,. A calibration procedure to improve global rice yield simulations with EPIC. Ecol Model. 2014; 128, 139.

36. Thorp KR, DeJonge KC, Kaleita AL, Batchelor WD, and Paz JO. Methodology for the use of DSSAT models for precision agriculture decision support. Comput Electron Agric. 2008;64:276–285. doi:10.1016/j.compag.2008.05.022

